# EPIC: An open community challenge for sequence-based prediction of transcription initiation in five non-model metazoans

**DOI:** 10.64898/2026.09.16.752171

**Authors:** Ilya E. Vorontsov, Nikita Gryzunov, Bayley R. McDonald, Vasily Kamenets, Timur Salimgareev, Ivan Kozin, The EPIC Consortium, Ashley M. Conard, Dmitry Penzar, Ivo Grosse, Vsevolod J. Makeev, Philipp Bucher, Ivan V. Kulakovskiy, Sascha H. Duttke

## Abstract

Predicting gene expression from DNA sequence is a central problem with critical biological and clinical implications. Recent sequence-to-function models are reported to achieve improved performance, yet it remains unclear how much of what they learn reflects genuine principles of eukaryotic transcription initiation and to what extent they are able to generalize beyond humans and other primary model species. A fair and blind benchmark has likewise been missing. Here we introduce EPIC, the <u>E</u>ukaryotic <u>P</u>romoter and transcription Initiation prediction <u>C</u>hallenge. Teams receive strand-specific, single-nucleotide-resolution initiation profiles for 80-95% of the genome and are asked to predict the remainder from DNA sequence alone. To level the field, the challenge relies on understudied animals spanning three phyla: octopus, oyster, milkweed bug, Indian meal moth, and shark. EPIC is open to everyone and closes on December 31, 2026. All teams clearing the dinucleotide precision baseline are invited to join the consortium authorship of the postchallenge publication, and winners are invited for personal authorship.

## 1. Introduction

Transcription of DNA into RNA is a first critical step in genome expression and fundamental to life. RNA is the template from which proteins are made, and often, RNAs are themselves key regulators or catalysts. Where transcription starts, and how efficiently, is encoded to a substantial degree in regulatory DNA via a multitude of short sequence motifs [1]. Understanding how these sequences encode transcription initiation is a central problem in biology and biomedicine, with critical implications for human health; most disease-associated mutations identified by genome-wide association studies fall outside protein-coding sequence and are enriched in regulatory DNA [2]. Being able to predict how a DNA sequence controls transcription initiation could therefore help us understand the molecular consequences of genetic variation, and ultimately identify mechanisms underlying human disease. The same regulatory code also shapes development and evolution. Changes in regulatory DNA can alter when, where, and how strongly genes are expressed, contributing to developmental innovations, evolutionary adaptation, and phenotypic diversity [3, 4]. Predictive modeling of transcription initiation from DNA sequence is thus of central importance.

Recent advances in machine learning provide an unprecedented opportunity to crack the “regulatory code”. Diverse base-pair-resolution models to predict initiation from sequence and recover interpretable promoter rules in the human genome have been developed [5-7], some across many modalities [8]. Genome language models trained across domains of life take a different route, learning sequence representation and conservation to predict function from sequence [9, 10]. However, where we are at in predicting transcription initiation from sequence is less clear. Conclusions about relative performance of a model depend heavily on how the evaluation is set up; results obtained on curated promoter or data subsets overestimate performance [11, 12], and whether a model actually learned principles that reflect biology or came to the right conclusion for the wrong reason is rarely tested [13, 14]. The <u>E</u>ukaryotic <u>P</u>romoter and transcription Initiation prediction <u>C</u>hallenge asks a simple but ambitious question: How well can we predict transcription initiation directly from DNA sequence?

EPIC follows those lessons and tries to address this need directly. Scoring is genome-wide over the whole eligible sequence rather than over a curated dataset, and results are evaluated against an unbiased, biological ground truth currently unknown to all participants, revealed at the end of the challenge. The DREAM [15] and CAGI [16] challenges established the format in regulatory genomics, and the recent IBIS challenge applied it to transcription factor binding specificities [17]. EPIC applies the same design to transcription initiation.

EPIC asks a deliberately narrow question. Given the genome sequence of five little-studied animals, including shark, oyster and octopus, and transcription initiation data for 80-95% of that genome, how accurately can models predict initiation for the remainder (Fig.1)? The main goal is to assemble a community-driven, open benchmarking suite for DNA sequence-to-transcription initiation prediction models. EPIC will further provide insights into how far the DNA syntax orchestrating transcription initiation is shared across animals and learned by models. Through these efforts, we may also learn where promoter architecture and initiation mechanisms are conserved and where they are plastic across the tree of life, and to what extent models learn transferable biological principles.

EPIC is enabled by capped small RNA sequencing (csRNA-seq), which captures active RNA polymerase II transcription initiation events at single-nucleotide resolution [18, 19]. Unlike approaches such as CAGE or 5′ scRNA-seq, which primarily measure stable, steady-state RNA 5′ ends, csRNA-seq directly captures active transcription initiation. It therefore detects not only stable protein-coding and non-coding genes, but also transient enhancer, promoter-antisense, and primary microRNA transcripts that are often missed by other approaches. This advance is critical: steady-state RNA 5′ ends reflect the combined effects of transcription initiation, elongation, and RNA degradation. By contrast, csRNA-seq provides a direct readout of transcription initiation itself, providing a more stringent test of sequence-to-function models and an opportunity to discover how the regulatory code of the genome controls transcription.

## 2. Challenge design

### 2.1 Experimental data

EPIC uses unpublished, strand-specific, single-nucleotide-resolution initiation csRNA-seq data from five evolutionarily diverse and largely understudied animals: two insects from different orders, an octopus, the Pacific oyster, and a cartilaginous shark (Fig. 1). The data are intentionally minimally processed. Reads are aligned to the respective species’ genomes and filtered for unique mapping, after which the 5′ ends of reads are tallied to generate strand-separated, genome-wide, single-nucleotide read-count profiles. No peak calling, cluster merging, or smoothing is applied [20]. The prediction target is the relative number and position of RNA polymerase II transcription initiation events with single-nucleotide and strand-specific resolution. EPIC does not target tRNAs or other small RNAs transcribed by RNA polymerase III, which are governed by distinct regulatory mechanisms and not captured by csRNA-seq. Highly abundant processed microRNAs and other small RNAs may be encountered. These can typically be identified and removed using matched small-RNA input data [19]. Small-RNA data are provided but for simplicity, and because such occurrences are rare, not necessary for this challenge: <u>EPIC does not use this additional small-RNA filtering step for preprocessing or scoring</u>. Participants, however, may use the small-RNA data to test and improve model performance.

**Figure 1.**
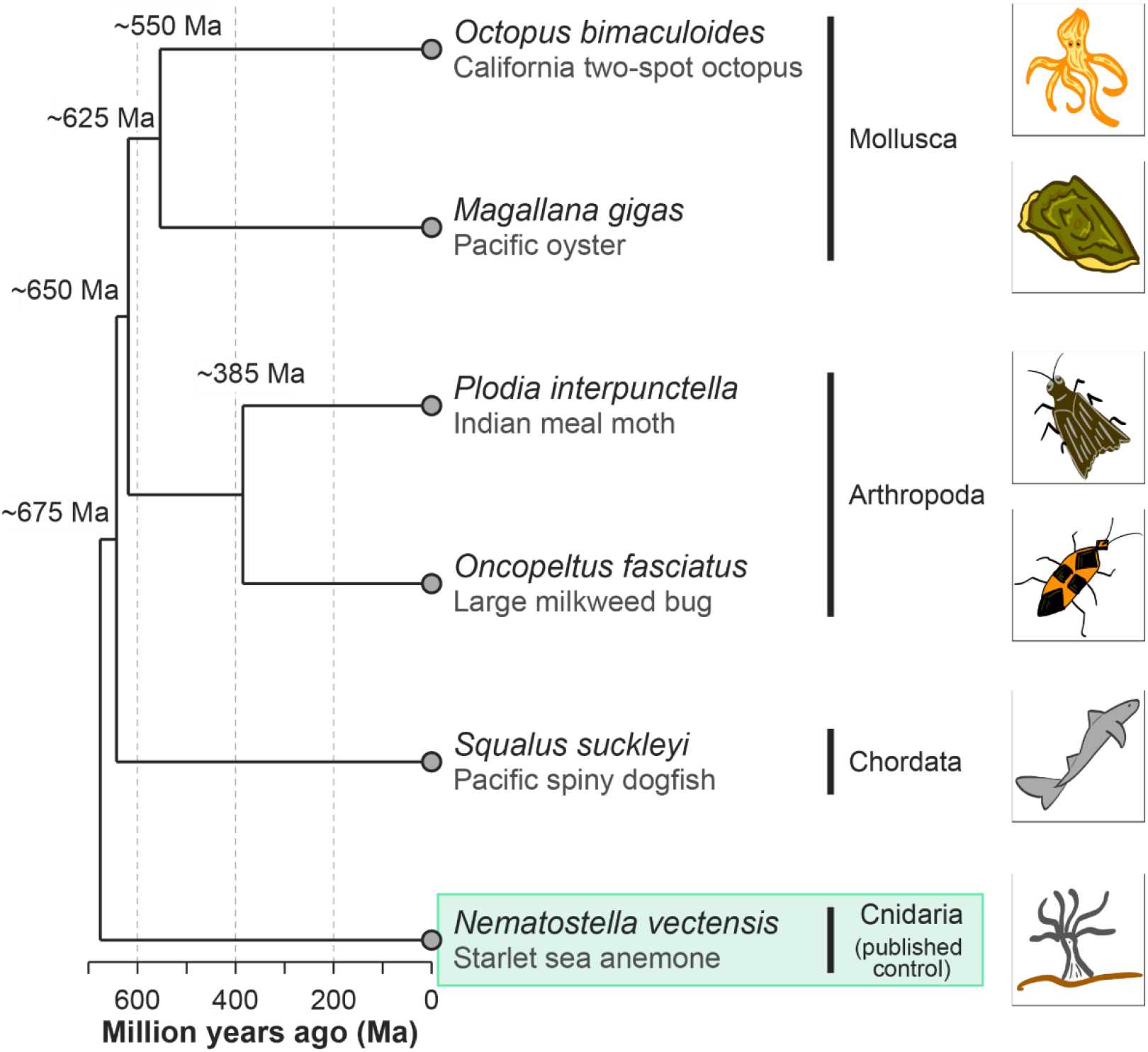
| EPIC - a community benchmark for predicting transcription initiation from sequence. The EPIC species span approximately 675 million years of animal evolution and represent three phyla. Approximate divergence times are shown at each node. Together, these species allow testing how well models generalize across genomes with very different composition and evolutionary history, rather than to a single well-studied species. The cnidarian *Nematostella vectensis* (starlet sea anemone; green box), whose transcription initiation profile is published, can be used for developing and validating pipelines against the challenge scoring before the sealed data are released.

### 2.2 Species and genomes

The benchmark spans three phyla (Fig.1); Assemblies and RepeatMasker annotations were taken from the UCSC GenArk hubs [21] so that every participant works from an identical reference. Genome sizes, contig counts, effective post-masking sizes, GC content, class balance, and train/test base-pair totals are distributed with the data package and are reproduced on the challenge website.

### 2.3 Train and test partition

For smaller genomes, approximately 80% of the eligible sequence of each genome is released for training, and approximately 20% GC-matched sequence is held out for evaluation. For the large genomes of *Octopus bimaculoides* and *Squalus suckleyi* we use 90%:10% and 95%:5% splits, respectively, to keep the submission file size manageable. Specifically,

1. Repeat-overlapping regions were removed, and remaining intervals shorter than 100 bp were discarded. Scoring uses only these repeat-free intervals, which are provided to participants as bed files.
2. Contigs are assigned to the training or the test set so that GC composition, computed on repeat-free sequence only, is balanced between the sets. GC content is the strongest global confounder for initiation prediction, and matching it prevents teams from being rewarded for learning the compositional idiosyncrasies of the test contigs.
3. The test set is then split into two equal parts, one used during the Leaderboard stage (continuously updated online) and the other assigned for the final evaluation.

Training data are released as gzipped BED files of per-position read counts (contig, start, end, id, readcount, strand), one file per replicate, in (1) a repeat-filtered version that matches how the test data will be scored and, as it might be beneficial to include repetitive regions in training, (2) an unfiltered version covering all positions of the training contigs.

For model training, per-replicate and pooled-replicate repeat-filtered tracks (gzipped bed and gzipped plain text) are provided together with the corresponding genome FASTA, its index, and the assembly report.

The target set of repeat-free genome regions for the test contigs is provided, but the experimental signal on those is withheld until the end of the challenge. For the test data, only the pooled-replicate track will be used for scoring and ranking the challenge solutions, although individual replicates might be used in the post-challenge assessment.

### 2.4 Prediction task and submission format

For each species, participants receive an ordered, strand-specific list of eligible genomic positions in BED format, for which they must predict the initiation strength. A submission is a plain gzipped text file with one prediction per line in template order (all plus strand predictions followed by all minus strand predictions; the participants should follow the provided templates carefully). Each line must contain a single non-negative real value in [0, 1] with up to five decimal places. One prediction file must be submitted per species; only gzipped .txt files which exactly match the template will be accepted. An offline validator script is provided to help participants check their submissions before uploading them. BED-formatted prediction files are not accepted. The same submitted predictions are used to evaluate both challenge tasks (see next section). Thus, a single prediction track must simultaneously distinguish initiating from non-initiating positions and accurately rank initiation strength among positions that are actively initiating.

### 2.5 Evaluation and ranking

Two tasks are scored per species:

- Classification (binary). Positions with non-zero transcription initiation counts are positives, and positions with no reads are negatives. Performance is measured as the area under the precision-recall curve (average precision). We adopted this measure as positives are a *small* fraction (<0.3%) of eligible strand-specific positions.
- Regression (quantitative). Restricted to the ground truth positives, the Spearman correlation between predicted and observed initiation levels (‘dense’ ranks).

Per-species scores from the two tasks are combined by log-rank aggregation, and species-level ranks are aggregated the same way into an overall ranking [17]. Gold, silver, and bronze medals are awarded on this basis for overall performance across species. Only a random half of the test labels contribute to the online leaderboard, which limits the gain available from fitting the leaderboard itself. Ground-truth profiles for test contigs will be announced with the winners at the end of the challenge. Clearing the simple basic dinucleotide predictor baseline threshold (Fig. 2) is required for joining the EPIC consortium.

**Figure 2.**
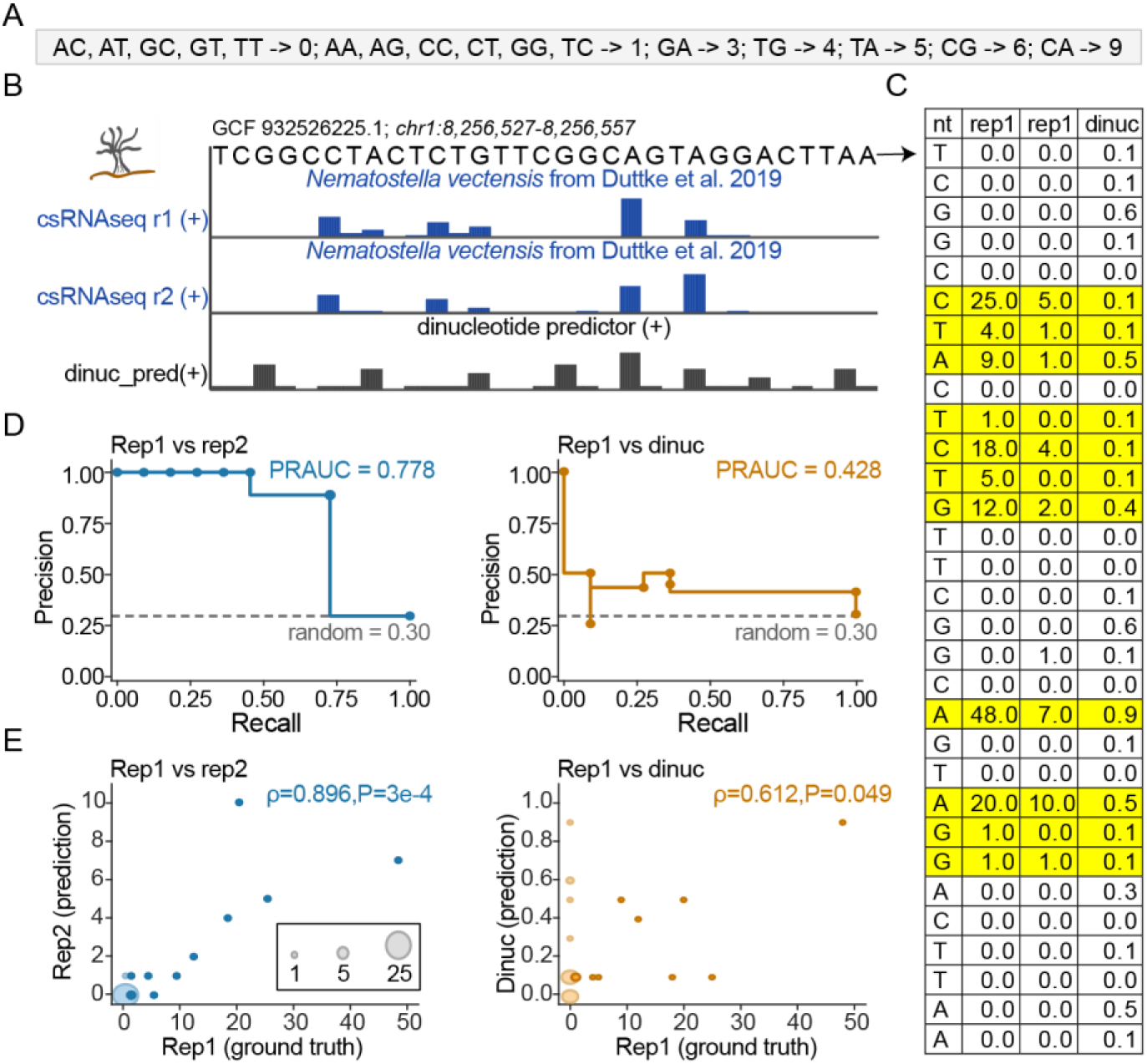
| Example of a simple dinucleotide predictor model. A.Base-line predictor scores: Initiation numbers were assigned to each dinucleotide based on input data. **B**. UCSC genome browser view of the challenge data and base-line predictions. **C**. Numerical data for the tracks shown in B. In yellow, positions that are used for computing correlation measures. **D**. AUPRC (Area Under the Precision-Recall Curve) for replicate and dinucleotide baseline predictor. Replicate 1 serves as ground truth. All positions were used for this analysis. **E**. Correlation of predictors with ground truth. Only yellow-marked positions in C (non-zero in ground truth) are used for this analysis.

### 2.6 Rules, permitted resources, and reproducibility

The goal of this challenge is clean sequence-to-signal comparison. Only the genome sequences and the EPIC-provided transcription initiation tracks may be used. Existing annotations, orthology relationships, expression data, other experimental tracks, or any other external data are not permitted. The exception is pretrained genomic models, such as (but not limited to) AlphaGenome [8] or Evo 2 [9] published in some form prior to the challenge start date, to allow comparison of these models against models trained from scratch.

Teams of one to ten members register through GitHub. A designated spokesperson must submit all predictions for the team. Every team willing to join the EPIC consortium must submit a method write-up accompanying the final submission together with training and scoring code sufficient to reproduce the submitted predictions. A leaderboard is live for the duration of the challenge; the final ranking will be made public after the deadline. Organizers are ineligible for medals.

To provide teams with the option of validating their pipelines before submitting, scoring scripts together with a published csRNA-seq dataset for the sea anemone *Nematostella vectensis* [19], processed identically to the challenge data, are provided.

## 3. Timeline and participation

The submission deadline is December 31, 2026. Registration, data access, and submission are handled at https://epic.autosome.org. Technical questions are answered in the EPIC Telegram group linked from the site.

## 4. Anticipated outcomes

We expect EPIC to produce:

- **A ranking of approaches** using identical, blind data; the first head-to-head comparison of sequence-based initiation prediction across phylogenetically distant animals.
- **A calibrated estimate of where the field stands** in predicting initiation from sequence.
- **A test of transfer**. Whether large pretrained genomic models retain useful information about initiation in genomes unlike their training data, and how this information transfer compares to compact models trained from scratch on the challenge data.
- **A peek into what the models have learned**. The core initiation machinery is broadly conserved across animals, while parts of the sequence grammar around it are not. A model that has captured general principles of initiation should perform consistently across species; one that rides on genome-specific composition should not. Five distant species, together with post-challenge analyses of promoter architecture, replicate agreement, and direct cross-species transfer, should provide an opportunity to disentangle these.
- **A persistent benchmark**. Data, templates, scoring code, and submitted predictions will be released alongside the post-challenge publication, so that later methods can be evaluated on the same terms.

## 5. Known limitations of the benchmark

Several design decisions constrain what EPIC can measure, and we state them explicitly.

- Repeat masking removes transposon-derived promoters from both training and scoring, so the benchmark deliberately scores the more tractable part of the genome.
- A single read creates a positive, so AUPRC is partly a function of species and library complexity. Thus, the resulting numerical values have different scales and are not directly comparable across species.
- Each species is represented by a single sample. Nothing in the benchmark tests condition-dependent or tissue-specific initiation.
- GC matching is performed at the level of whole contigs and therefore balances the marginal composition of the training and test sets rather than local composition around initiation sites.

## 6. Call for participation

Anyone is invited and can participate! No prior work or knowledge of transcription initiation is required. Data are provided in a standard and convenient format. None of the five species selected for EPIC have been previously studied for this purpose, so no participant starts with valuable species-specific knowledge that others lack. The task is self-contained: sequence in, per-position score out. Single participants and small teams are welcome alongside groups of up to ten. Registration and data access are at https://epic.autosome.org.

### Consortium authorship

Every team clearing the baseline and providing the method write-up is eligible to join the EPIC Consortium and will be listed as consortium authors on the post-challenge publication. The challenge winners are invited as named individual co-authors on the EPIC challenge manuscript.

### Data usage policy

Training data, genome references, submission templates, and scoring scripts are distributed through the challenge website upon registration. The training and test data derive from unpublished experiments, and participants are asked not to redistribute the data outside this challenge without permission from the organizers. Teams intending to publish a method developed on the EPIC benchmark are asked to wait until the challenge and data preprints are posted. The test-set ground truth will be released upon the challenge completion and verification of the winning solutions.

## Data and code availability

Scoring and offline validation scripts are available at https://github.com/autosomeru/EPIC_scoring

The EPIC data are available at Zenodo: https://doi.org/10.5281/zenodo.22285753

## Author contributions

Challenge conception and organization: P.B., I.V.K., S.H.D. Steering committee: I.G., V.J.M. Experimental data production and curation: B.R.M., I.V.K., S.H.D. Benchmarking infrastructure and software: I.E.V., N.G., A.M.C., T.S., I.K., D.P. All authors contributed to the challenge design and to this manuscript.

## Funding and acknowledgements

We thank the IT Group of the Institute of Computer Science at Martin Luther University Halle-Wittenberg for computational resources and, personally, Maximilian Biermann, for valuable technical support. S.H.D. is supported in part by the Microsoft Research Fellowship “Generative models for regulatory genomics” and NIH grant R00GM135515. I.V.K., I.E.V., I.K., and D.P. are supported by assignment 125091010189-3 to I.V.K.

